# Causal inference during closed-loop navigation: parsing of self- and object-motion

**DOI:** 10.1101/2023.01.27.525974

**Authors:** Jean-Paul Noel, Johannes Bill, Haoran Ding, John Vastola, Gregory C. DeAngelis, Dora E. Angelaki, Jan Drugowitsch

## Abstract

A key computation in building adaptive internal models of the external world is to ascribe sensory signals to their likely cause(s), a process of Bayesian Causal Inference (CI). CI is well studied within the framework of two-alternative forced-choice tasks, but less well understood within the cadre of naturalistic action-perception loops. Here, we examine the process of disambiguating retinal motion caused by self- and/or object-motion during closed-loop navigation. First, we derive a normative account specifying how observers ought to intercept hidden and moving targets given their belief over (i) whether retinal motion was caused by the target moving, and (ii) if so, with what velocity. Next, in line with the modeling results, we show that humans report targets as stationary and steer toward their initial rather than final position more often when they are themselves moving, suggesting a misattribution of object-motion to the self. Further, we predict that observers should misattribute retinal motion more often: (i) during passive rather than active self-motion (given the lack of an efference copy informing self-motion estimates in the former), and (ii) when targets are presented eccentrically rather than centrally (given that lateral self-motion flow vectors are larger at eccentric locations during forward self-motion). Results confirm both of these predictions. Lastly, analysis of eye-movements show that, while initial saccades toward targets are largely accurate regardless of the self-motion condition, subsequent gaze pursuit was modulated by target velocity during object-only motion, but not during concurrent object- and self-motion. These results demonstrate CI within action-perception loops, and suggest a protracted temporal unfolding of the computations characterizing CI.

## 1. Introduction

We do not directly access environmental objects and events. Instead, our biological sensors (i.e., retina, cochlea, etc.) detect noisy, incomplete, and often ambiguous sensory signals. In turn, our brains ought to leverage these signals to build adaptive internal models of the external world [1]. A key step in building these models is to ascribe sensory signals to their likely (and hidden) cause(s), a process of Bayesian causal inference (CI; [2–4]).

CI has been well studied in human psychophysics, and a growing body of literature is starting to elucidate the neural mechanisms underpinning this computation, both in humans [5–10] and animal models [11–13]. Namely, whether it is localizing audio-visual stimuli [2, 5, 11], estimating heading [6, 12], perceiving motion relations in visual scenes [9, 10], or inferring the location of one’s own body based on visual, tactile, and proprioceptive cues [7, 8, 13], humans and non-human primates behave as if they hold and combine (sometimes optimally) multiple interpretations of the same sensory stimuli. From a neural standpoint, CI appears to be similarly subserved by a cascade of concurrent interpretations; sensory areas may respond to their modality of preference, intermediate “associative” nodes may always combine cues, and finally higher-order fronto-parietal areas may flexibly change their responses based on the causal structure inferred to generate sensory observations [4, 14, 15] (see [16, 17] for similar findings across time, from segregation to integration to causal inference). If and how this inferred causal structure subsequently and dynamically biases lower-level sensory representations is unknown (but see [18]).

More broadly, the study of CI has heavily relied on static tasks defined by binary behavioral outcomes and artificially segregating periods of action from those of perception (see [19] for similar arguments). These tasks are not only a far cry from the complex, closed-loop, and continuous-time challenges that exist in human behavior, but may also limit and color our understanding of CI. For instance, while feedforward-only mechanistic models of CI may account for decisions during two-alternative forced choice tasks [20], our brains are (i) decidedly recurrent, and (ii) largely dictate the timing, content, and relative resolution of sensory input via motor output (i.e., active sensing). The focus on open-loop tasks when studying CI also limits our ability to bridge between CI and other foundational theories of brain function – particularly those derived from reinforcement learning, which are best expressed within closed loops.

Here, we take a first step toward understanding CI under closed-loop active sensing. Human observers are tasked with navigating in virtual reality to the location of a briefly flashed target, much like the “blinking of a firefly” (see [21–25]). The virtual scene is composed solely of flickering ground plane elements, and thus, when observers move by deflecting a joystick (velocity control), the ground plane elements create optic flow vectors. Observers continuously integrate this velocity signal into an evolving estimate of position [26]. Importantly, in the current instantiation of the task, the target may itself move (i.e., object-motion). Thus, in the case of concurrent self- and object-motion, observers must parse the total retinal motion into components caused by self-motion and/or those caused by object-motion, a process of CI.

First, we derive a normative account specifying how observers ought to intercept hidden and moving targets given their belief over (i) whether retinal motion was (at least partially) caused by the target moving, and (ii) if so, at what velocity. Then, we demonstrate that humans’ explicit reports and steering behavior concord with the model’s predictions. Further, we support the claim that humans misattribute flow vectors caused by object-motion to their self-motion by showing that these mis-attributions are larger when i) self-motion is passive (i.e., lacking an efference copy) and when ii) objects are presented eccentrically (such that object and self-motion vectors are congruent in direction). Lastly, via eye-movement analyses we show a behavioral unfolding during CI. Early saccades are largely accurate and directed toward the last visible location of the target. Thus, during this time period eye-movements behave as if participants perceived targets as moving. Subsequent gaze pursuit of the invisible target is accurate when there is no concurrent self-motion, but is consistent with the target being perceived as stationary during concurrent self- and object-motion. Together, the results demonstrate CI in attributing retinal motion to self- and object-motion during closed-loop goal-directed navigation. This opens a new avenue of study wherein we may attempt to understand not only the perceptual networks underpinning CI, but the joint sensorimotor ones.

## 2. Results

### 2.1 A normative model of intercept behavior during visual path integration

We identify behavioral signatures of CI during path integration toward a (potentially) moving target by formulating a normative model (**Fig. 1**). We take the example of a briefly visible target moving at a constant, potentially zero, speed (i.e., no acceleration) and with a constant direction along a lateral plane (i.e., side-to-side, **Fig. 1A**). We model concurrent self- and object-motion (vs. object-motion only) by increasing the noise associated with target observations (these being corrupted by a concurrent flow field). The model first forms a Bayesian belief over the target’s location and velocity given the brief observation period (i.e., period over which the target is visible). It then uses this belief to steer and intercept the target. For simplicity, within this work we only consider linear trajectories wherein the model selects a direction and duration over which to travel (**Fig. 1A** middle panel). We expect this trajectory to well-approximate the steering endpoints of the continuously controlled steering trajectories of our human participants.

**Figure 1:**
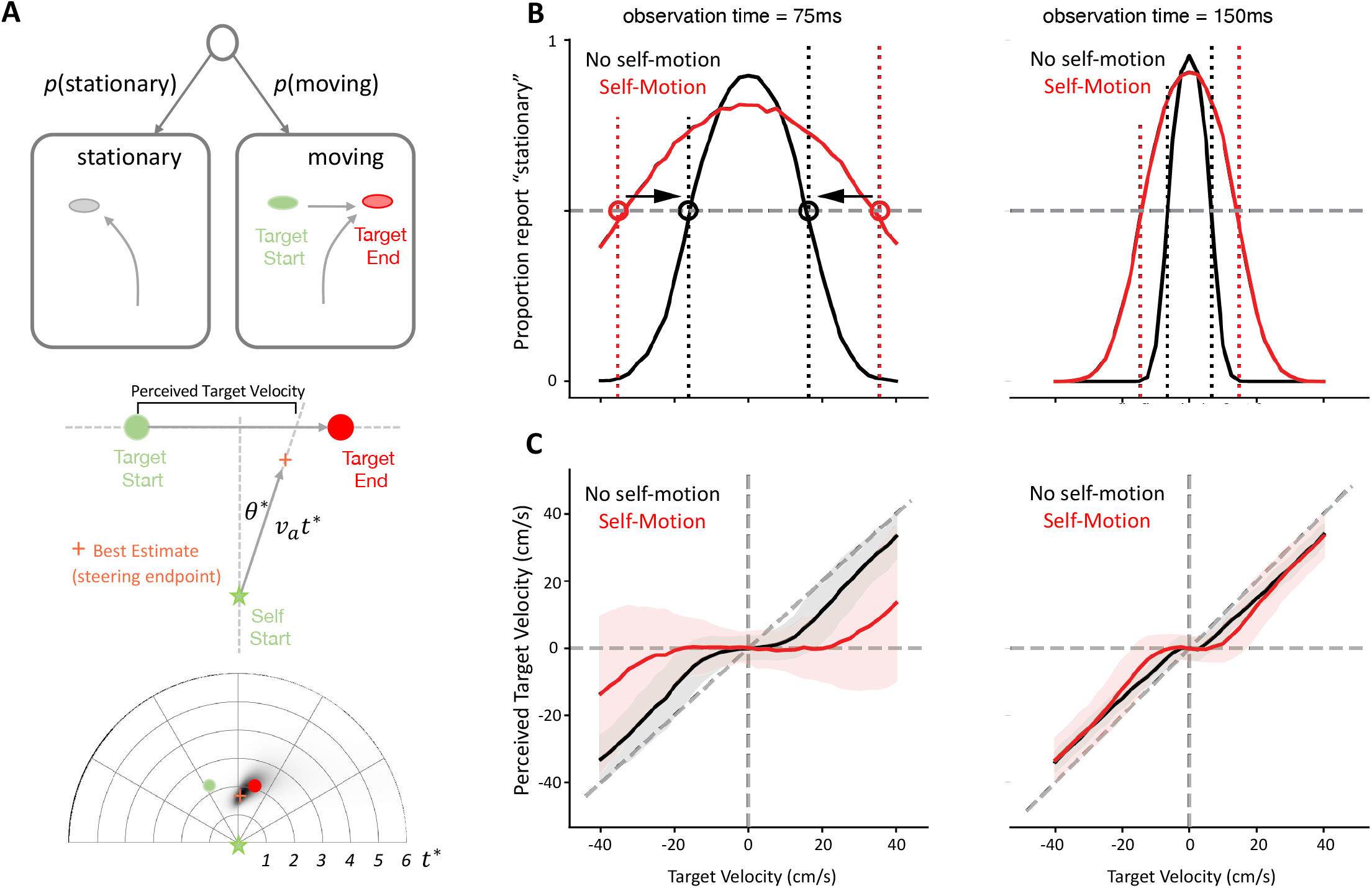
Predicted behavioral signatures of causal inference when navigating to intercept a briefly observed moving target. A. Task and trajectories of a normative model. We derived the normative strategy for intercepting a briefly presented target that provided the model with an uncertain estimate of the target velocity and whether the target was moving at all. Top: the model first estimates whether the target is stationary or moving. Middle: we assumed that the model aimed to intercept the target by traveling along a straight line at a certain angle (*θ*^*^) and for a certain distance (velocity x duration, *v*_*a*_*t*^*^). To compensate for the model overshooting or undershooting the target (given uncertainty in path integration, see bottom panel), we assessed the model’s perceived velocity by computing when the model’s trajectory intercepted the target’s path. Middle panel shows a schematized example, while bottom panel shows an example simulation including the best estimate endpoint (orange) and full posterior (shades of black) **B. Stationary reports**. The model perceived the target as stationary if its noisy velocity estimate fell below a velocity threshold (dashed vertical lines). Noise in the velocity estimates resulted in a bell-shaped fraction of stationary reports when plotted over the target’s true velocities. For a Bayesian decision strategy, and in contrast to simpler heuristics, the velocity thresholds increased for larger observation noise induced by self-motion (red vs. black) and for shorter observation times (left vs. right). The bell-shaped curves widened accordingly, and the point at which they intersected the 0.5 proportion (grey dashed line) changed. We highlight this threshold change here for the self-motion vs. no self-motion case by the circles and associated black arrows. **C. Perceived target velocities**. Causal inference causes the perceived velocities to be biased toward zero for small target velocities, and to approach true target velocities (down-weighted by the slow-velocity prior) for larger target velocities. This bias increases for larger observation noise (red vs. black) and shorter observation periods (left vs. right). **B** and **C** show mean (lines) and SD (shaded area, **C** only) across 1000 simulated trajectories for each target velocity, ranging from −40 to 40 cm/s in steps of 2.5cm/s.

On each trial, before making any observation, the model assumes the target to be stationary with a certain probability, and to move otherwise (i.e., a prior for stationarity). If moving, it assumes that slower targets are more likely than faster ones (i.e., a slow-velocity prior, [27, 28]). When the target is rendered visible, the model gathers noisy observations of the target’s location (e.g., noisy percepts in each video frame, potentially corrupted by a concurrent flow field) that it uses to infer whether the target is stationary or moving. And if moving, with what velocity. Once it has formed these target motion estimates, it uses a simple steering policy in which it moves a certain distance (determined by a velocity and optimal stopping time) along a straight-line trajectory to best intercept the target (**Fig. 1A**, see *Methods* and *Supplementary Information* for details).

Simulating the model across multiple trials led to two predictions. The first prediction is that whether the target is perceived as stationary or moving should depend on the target’s actual velocity, the observation time, and observation noise (i.e., “no self-motion” vs. “self-motion”, respectively in black and red; **Fig. 1B**). Noisy observations during concurrent self-motion imply that a moving target might be mistaken for a stationary one, in particular if this is a-priori deemed likely. Our model defaults to such a stationary target percept as long as the target velocity estimate remains below a specific velocity threshold (**Fig. 1B**, dashed vertical lines). Noise in the target velocity estimates thus leads to a bell-shaped relationship between the probability of a stationary target percept and the target’s actual velocity. This relationship is modulated by observation time and observation noise magnitude: both larger observation noise magnitudes and shorter observation times require stronger evidence to perceive a target as moving (**Fig. 1B**, black vs. red and left vs. right). Our model implements this by increasing the threshold on the target velocity estimate. This change in threshold provides a test for whether observer (i.e., human) behavior is truly Bayesian, in the sense that it is sensitive to changes in the observation quality [29]. If the moving/stationary decision is implemented by a fixed heuristic, rather than Bayesian velocity threshold, then the target velocity that leads to a probability of stationary percepts of 0.5 would not shift with a change in observation noise or time (in contrast to arrows shown in **Fig. 1B**).

The second prediction relates the target’s true velocities to those perceived by the model. In each simulated trial, the model uses the target motion estimate to steer along a straight trajectory that maximizes the likelihood of intercepting the target, while accounting for standard path integration properties (e.g., increasing location uncertainty with distance traveled, see *Methods* and [21]). To estimate our model’s perceived target velocity, we intersected a straight line connecting steering start and endpoints with the line that the target moved along (which was the same across trials, **Fig. 1A**). This procedure compensates for the steering endpoint occasionally overshooting or undershooting the target’s location, given noisy path integration and speed priors (see [21] and below). Plotting these perceived velocities over true target velocities revealed characteristic S-shaped curves (**Fig. 1C**). For large target velocities, the perceived velocities linearly increase with target velocities, but consistently underestimate them due to the slow speed prior. Smaller target velocities reveal signatures of CI: as small target velocities occasionally make the target appear stationary, the perceived velocities are further biased toward zero. Both larger observation noise magnitudes and shorter observation times expand the range of true target velocities for which the perceived velocities are biased toward zero (**Fig. 1C**, black vs. red and left vs. right), in line with the stationarity reports (**Fig. 1B**).

In summary, therefore, according to the normative model, if concurrent self- and object-motion leads to increased observation noise (vis-à-vis a condition with no concurrent self-motion), then we ought to expect (i) a larger velocity range over which targets are reported as stationary, and (ii) a characteristic S-shaped curve where observers more readily navigate toward the starting rather than ending location of targets.

### 2.2 Human observers perform causal inference when navigating during concurrent object-motion

We test predictions of the model by having human observers (n = 11) navigate by path integration to the location of targets that could either be stationary (i.e., no object velocity) or moving with different velocities relative to the virtual environment (**Fig. 2A**, dashed black line denotes object velocity). As in the model, targets moved at a constant speed (range from 0 to 40cm/s) and with a constant lateral direction (i.e., leftward or rightward, if moving). Further, at the end of each trial participants explicitly reported whether they perceived the target as moving or not relative to the scene. Most importantly, in different blocks of trials the observers themselves could either be stationary (i.e., labeled “no self-motion” and requiring participants to maintain a linear velocity under 1cm/s for 1 second for targets to appear) or moving during the time period when the target was visible (i.e., labeled “self-motion” and requiring participants to maintain a linear velocity over 20cm/s for 1 second for targets to appear). The optic flow caused by self-motion introduces additional observation noise and requires the parsing of total flow vectors into self- and object-components.

**Figure 2.**
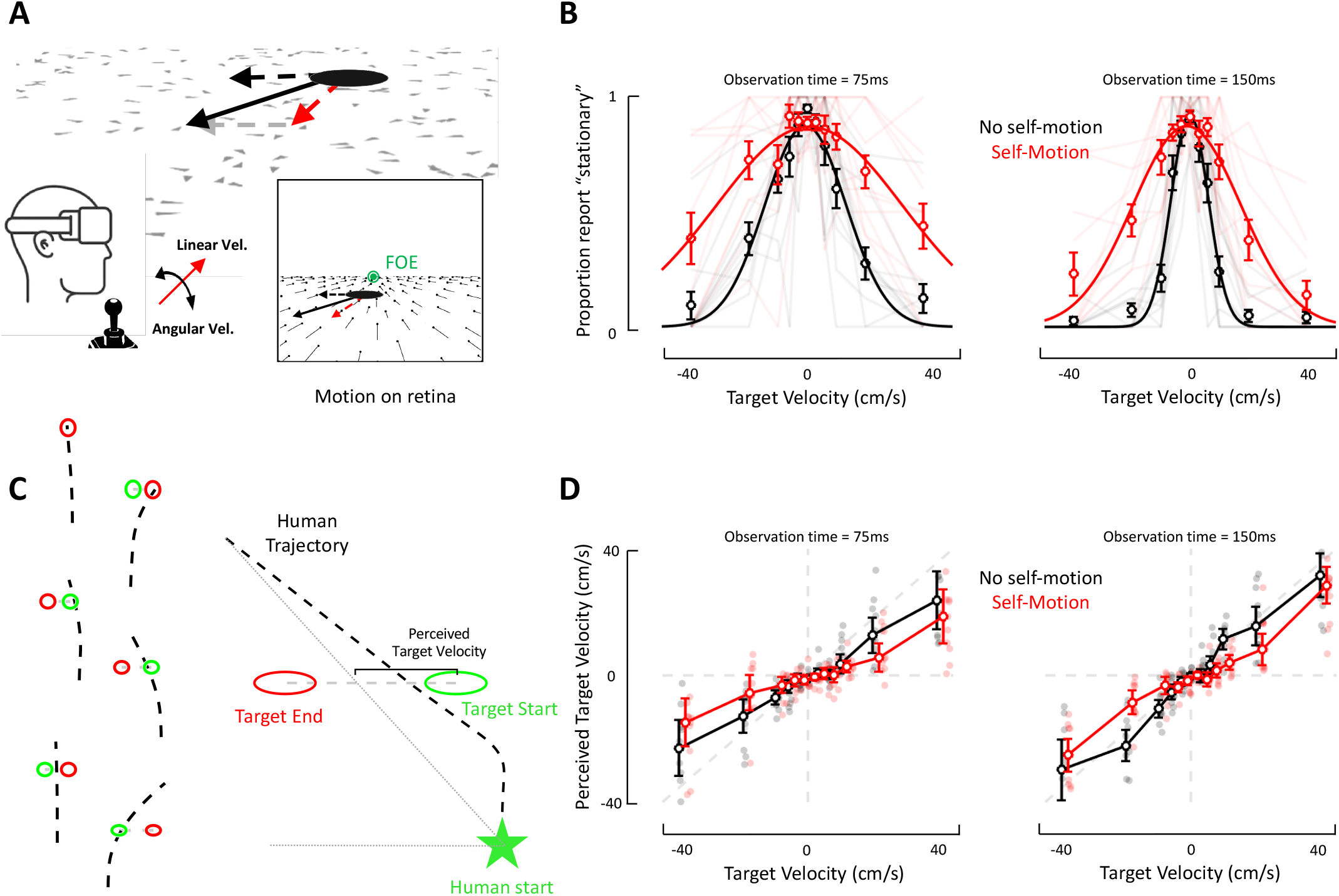
Intercepting a briefly visible moving target by path integration demonstrates features of causal inference. **A. Experimental protocol and setup.** Participants are placed in a virtual scene composed of intermittently flashing textural elements providing an optic flow signal when participants are moving. Subjects are in closed-loop, at all times being in control of their linear and angular velocity. When the target has an independent motion in the environment (dashed black line) and subjects move toward it (red lines), the retinal motion of the target (solid black line) is composed of both object- and self-motion components. Inset additionally shows flow vectors. FOE (in green) indicates the focus of expansion. **B. Stationary reports**. The proportion of trials in which participants reported the target as stationary (y-axis) is plotted as a function of target velocity (x-axis), whether the subject was moving (self-motion, red) or not (no self-motion, black) during target presentation, and target observation time (left vs. right panels).Circles denote means across subjects, and error bars represent +/− 1 standard error of the mean (S.E.M). Transparent lines in the background show data for individual subjects. **C. Example steering trajectories and quantification**. Left: Six example trials (arbitrarily staggered). Bird’s eye view of the target’s starting (green circle) and ending (red circle) location, as well as individual trajectories (dashed black). The distance between the start of the dashed black line (origin) and the target’s starting location is always 300cm. Right: To estimate the perceived target velocity, we computed for each trial the distance between the target’s initial location, and the location where the target’s trajectory intersects a straight line connecting the human’s starting and ending locations. This recapitulates the definition from **Fig. 1** and defines the lateral displacement of an individual trajectory above and beyond the starting location of the target, while also considering the depth overshooting. **D. Perceived target velocity**. The perceived target velocity (as quantified in **C**, y-axis)) is plotted as a function of actual target velocity (x-axis), whether the subject was moving (self-motion, red) or not (no self-motion, black) during target presentation, and target observation time (left vs. right panels). Circles denote means, and error bars represent +/− 1 S.E.M. Individual transparent dots in the background show data for individual subjects.

Explicit stationarity reports were well accounted for by Gaussian distributions (mean r^2^ = 0.74). As expected, participants most readily reported targets as stationary in the world when the object velocity was close to 0 (mean of Gaussian fits pooled across “self-motion” and “no self-motion” conditions; 75ms observation time: *μ* = −4.65 x 10^−4^; 150ms observation time: *μ* = 9.2 x 10^−3^), regardless of the target observation time (p = 0.47). Most importantly, as predicted by the model, during concurrent self- and object-motion (red, **Fig. 2B**) participants reported targets as stationary at increasing object velocities, both for lower (75ms; Gaussian standard deviation, mean ± s.e.m., no self-motion: 10.89cm/s ± 2.37cm/s; self-motion: 29.39cm/s ± 7.06cm/s; paired t-test, p = 0.021) and higher (150ms; no self-motion: 4.59cm/s ± 0.51cm/s; self-motion: 15.15cm/s ± 3.01cm/s; p = 0.0025) observation times. This suggests that participants putatively misattributed flow vectors caused by object-motion to self-motion.

We similarly analyzed the end points of participants’ steering behavior. This behavior was heterogenous, with participants frequently stopping at the target’s end location (**Fig. 2C**, top left and right; examples of stationary and moving targets), but also at times navigating to the starting and not ending target location (**Fig. 2C**, middle left) or navigating to some intermediary location (**Fig. 2C**, bottom left). Likewise, participants often overshot targets in depth (**Fig. 2C**, middle and bottom right, also see [19], radial response divided by radial target distance, no self-motion = 1.44 ± 0.08; self-motion = 1.32 ± 0.09; no self-motion vs. self-motion, t = 1.55, p = 0.15). To quantify this performance, for each trial we computed a perceived target velocity analogously to the modeling approach (**Fig. 1A** and **2C**). Results demonstrated that, on average, subjects underestimated the target’s velocity (linear fit to no self-motion condition, grand average slope = 0.82 ± 0.08; slope = 1 indicates no bias), and this effect was exacerbated during low observation times (**Fig. 2D**, slopes of linear fits, 75ms vs. 150ms observation times paired t-test, p = 0.05). Interestingly, when targets were presented during concurrent self-motion (**Fig. 2D**, red), intermediate target velocities were perceived (or at least steered toward) as if moving more slowly than when the same object velocity was presented in the absence of self-motion (**Fig. 2D**, red vs. black, paired ANOVA interaction term, 75ms observation time, p = 8.12 x 10^−5^; 150ms observation time, p = 1.53 x 10^−9^; Bonferroni-corrected p < 0.05 for 75ms observation time at −20cm/s, −6cm/s and +10cms, and for 150ms observation time at −20cm/s, −10cm/s, and +10cm/s).

### 2.3 Steering behavior suggests a misattribution of flow vectors during concurrent self- and object-motion

To further test the claim that during concurrent object-motion and path integration participants may misattribute flow vectors related to the object and self, we can make two further qualitative predictions. First, we predict that the location of the target at its onset will affect how object- and self-motion derived flow vectors are misattributed. That is, early in trials, participants move forward, toward the target. During this forward self-motion, lateral flow vectors on the retina – i.e., those matching the direction of target movement – are essentially null straight-ahead and have much greater speed in the periphery (see **Fig. 2A** inset). Thus, we predict that flow vectors should be more readily misattributed under our current protocol for eccentric, rather than central, targets. Second, within the current closed-loop experiment, self-motion is not solely estimated based on optic flow signals, but also by an efference copy of hand movements (and thus joystick position, which drives virtual velocity). In turn, we can predict that if self-motion during object-motion were passive (i.e., no efference copy), the estimate of self-motion would be more uncertain, and thus putatively more amenable to misattribution between object- and self-motion cues. We test these predictions while focusing our analysis on (1) the 150ms target observation time given that participants are more accurate in this condition, and (2) steering behavior (vs. explicit reports) given that these are an implicit measure of CI not prone to response biases ([27]).

We test the first prediction by dividing the dataset into trials for which the target was presented centrally (i.e., within −5° and +5°; 48.5% of total dataset) or eccentrically (i.e., within −10° and −5°, or within +5° and +10°). As shown in **Fig. 3A**, during both central and eccentric presentations, targets were seemingly perceived as if moving less rapidly during self-motion rather than during no self-motion. To quantitatively ascertain whether flow vectors were more readily misattributed during eccentric rather than central presentations, for each condition (central vs. eccentric) and target velocity we computed the difference in perceived target velocity (self-motion minus no self-motion). This difference is illustrated in **Fig. 3B**, and statistical contrasts indicated that the impact of self-motion was greater for eccentric rather than central presentation for targets moving at −10cm/s and 20cm/s (red vs. black, paired ANOVA object-velocity x self-motion condition interaction term, p = 0.03, post-hoc comparisons at each velocity, p < 0.05 Bonferroni-corrected at −10cm/s and 20cm/s).

**Figure 3.**
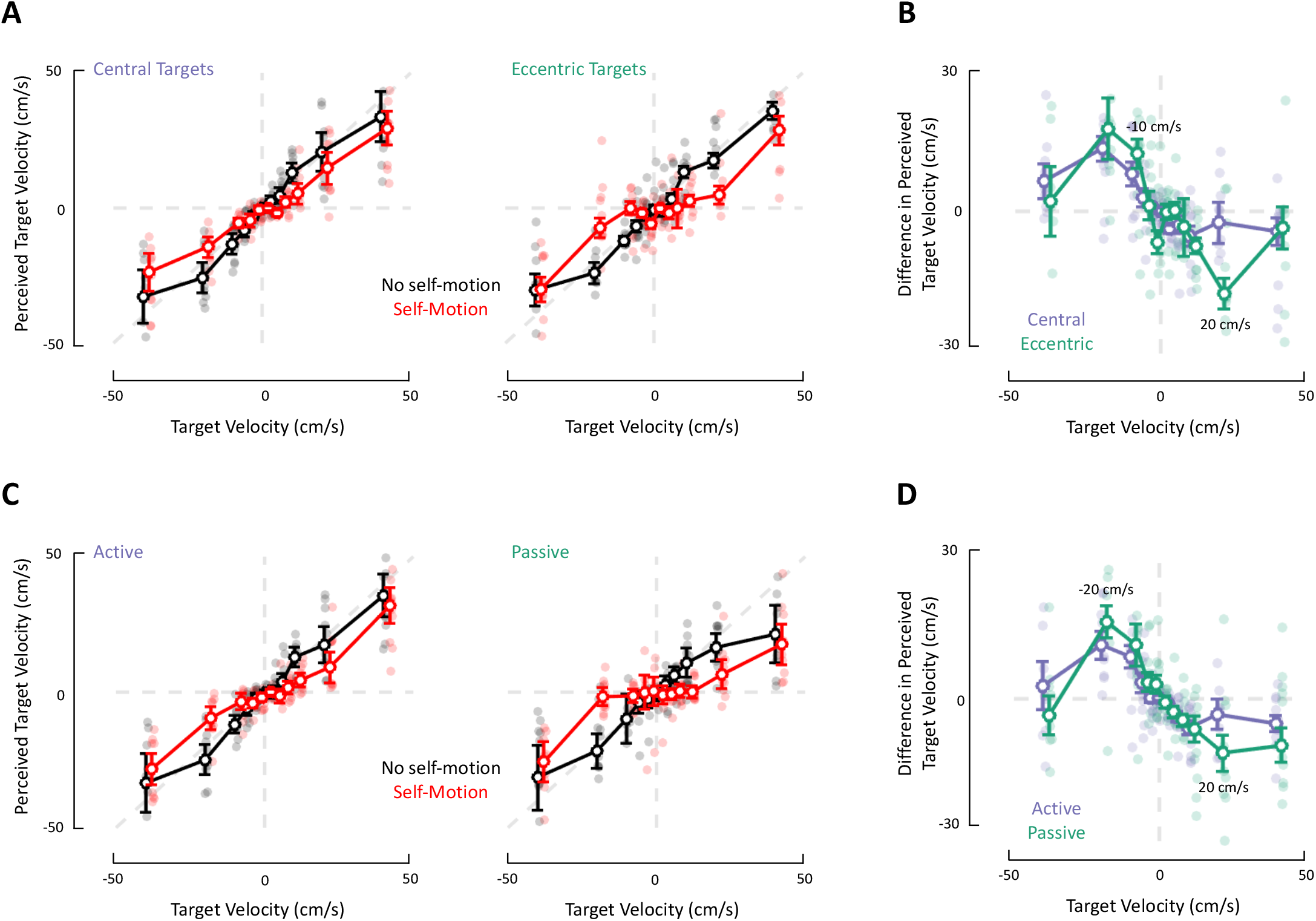
Misattribution of optic flow vectors during concurrent self- and object-motion is exacerbated for eccentric targets and during passive rather than active self-motion. **A. Effect of target location.** Perceived target velocity (y-axis, as estimated based on steering end-point) as a function of true target velocity (x-axis), whether participants were moving (red; self-motion) or not (black; no self-motion) while the target was visible, and whether targets at onset were central (left) or eccentric (right). Circles represent means, error bars denote +/− 1 S.E.M, and transparent dots in the background show data for individual subjects. **B. Impact of self-motion on perceived object-velocity as a function of target location**. Difference in perceived target velocity as a function of self-motion condition (self-motion - no self-motion; or red – black from **Fig. 3A**), and whether targets were central (purple) or eccentric (green) at onset. Circles indicate means, error bars denote +/− 1 S.E.M, and transparent dots in the background show data from individual subjects. **C. Active vs. passive self-motion**. Similar to **A**, but panels are separated as a function of whether self-motion during target presentation was active (i.e., closed-loop; left, data is reproduced from **Fig. 2D**, right) or passive (i.e., open-loop; right). **D. Impact of active vs. passive self-motion on perceived object-velocity**. Similar to **B**, but separated according to whether self-motion during target presentation was active (purple) or passive (green).

To test the second prediction, we conducted an additional experiment (same participants, see *Methods*) wherein participants’ linear velocity during target presentation (and only during this time-period) was under experimental control and was varied from trial-to-trial (either 0cm/s or a Gaussian profile ramping over 1 second to a maximum velocity of 200cm/s). In turn, we contrast the impact of self-motion on perceived target velocity either during closed-loop, active navigation (**Fig. 3C**, left, same data as in **Fig. 2D**, 150ms observation time) and during passive self-motion (**Fig. 3C**, right). As above, for both active and passive conditions we compute the difference in perceived target velocity between self-motion and no self-motion conditions. Results show that when targets moved at a velocity of −20cm/s and 20cm/s (**Fig. 3D**; ANOVA object-velocity x active vs. passive interaction term, p = 0.04, post-hoc comparisons, p < 0.05, Bonferroni-corrected at −20cm/s and 20cm/s), the misattribution of flow vectors during self-motion was greater during passive rather than active self-motion. Together, these results confirm our qualitative predictions, suggesting that optic flow vectors were misattributed during concurrent self- and object-motion. Further, the increased misattribution during eccentric target presentation and during passive self-motion was specific for intermediate firefly velocities, precisely when internal models are less certain.

### 2.4 Eye movements suggest a temporal cascade from segregation to integration to causal inference

Lastly, we attempt to leverage the continuous-time nature of the task to gain insight into the timecourse of the computations supporting CI during closed-loop navigation. Namely, the above analyses were restricted to trajectory endpoints, the outcome of the computation. However, we may also use the eye movements occuring during and near the time of target presentation, which is the critical period during which the task-relevant inference ought to occur.

Eye movements were composed of a rapid, ballistic saccade (**Fig. 4A**, thin lines show example trials and thicker lines show mean values, arrows indicate the mean latency to saccade offset for eccentric targets) followed by a more protracted smooth pursuit (**Fig. 4A**, slow drift after arrow). In prior work, we have demonstrated that humans (and non-human primates) intuitively saccade to the visible target, and then track it via smooth pursuit, even when hidden ([22, 25]). Unlike in these prior studies, however, here the target itself moves, and the presentation times were half as long (300ms before vs. maximum of 150ms here). Thus, the saccade itself happens after target offset (**Fig. 4A**). In turn, saccades and gaze pursuit may aim toward the target onset position, its final visible offset location, an intermediate location, and/or track an evolving location.

**Figure 4.**
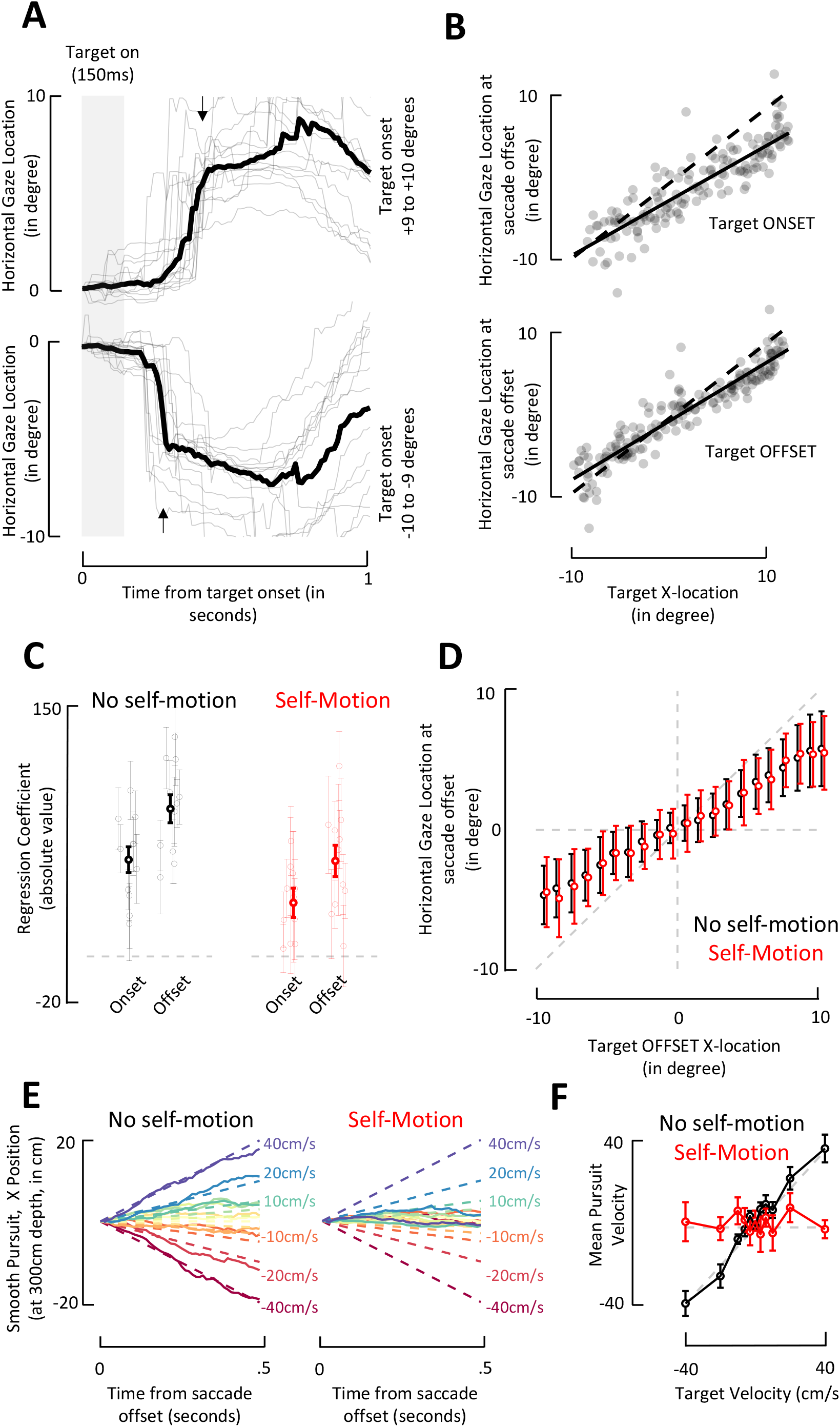
Saccade and smooth pursuit behavior during and following the target presentation show differential impact of self-motion. **A. Example lateral gaze position.** Top panel shows gaze (eye + head) direction along the lateral plane (azimuth) for targets far to the right (positive values; +9 to +10 degrees), while bottom panel shows gaze direction along the lateral plane for targets far to the left. Thin and transparent lines are examples (no self-motion condition), while the opaque and thicker line is the average. Arrows represent the mean latency of saccade. These representative data show a ballistic saccade (marked by the arrows) followed by smooth pursuit. The saccades happen after target offset (maximum target presentation time represented by gray box). **B. Saccade landing locations along the lateral plane for a representative subject**. Top; lateral gaze location (in degrees) at saccade offset as a function of the targets initial (onset) location (also in degrees). Bottom; as above, for target offset location (i.e., last visible location). Dashed line is the identity, while solid line is the best linear fit. Individual dots are trials. **C. Coefficient (in absolute value) for a multiple regression accounting for saccade offset location as a function of target onset and offset location**. This analysis was conducted both for the no self-motion (black) and self-motion (red) conditions. Thin and transparent error bars denote individual subjects and their standard error, while opaque and thicker error bars show the mean and s.e.m. Coefficients were larger for the offset than onset target location, both for the no self-motion and self-motion condition. **D. Lateral gaze location at saccade offset as a function of target offset**. Target offset locations were categorized in 21 bins (every degree, from −10 to +10 degrees) and gaze offset location was averaged for each subject within these bins. Dots show means across subjects, and error bars represent +/− 1 s.e.m. Dashed gray line is the unity-slope diagonal, demonstrating that subjects slightly undershot targets, but were largely accurate. **E. Baseline-corrected lateral gaze position after saccade offset during no self-motion (left) and self-motion (right) conditions**. The time-course of gaze (500ms after saccade offset) shows modulation as a function of the target velocity (gradient; from warm to cold colors; −40cm/s to +40cm/s) in the no self-motion condition, but no modulation during the self-motion condition. Solid lines are averages across all subjects. Dashed lines show the evolving location of targets. **F. Smooth pursuit velocity as a function of target velocity and self-motion condition**. During self-motion (red), gaze velocity (y-axis) was not modulated as a function of target velocity (x-axis). In contrast, gaze velocity did accurately track target velocity in the no self-motion condition (black). Dots show means across subjects, and error bars represent +/− 1 s.e.m. Dashed gray lines show y = 0 and y = x.

To examine saccades, we expressed gaze location along azimuth by adding eye-in-head orientation and head orientation. Similarly, we expressed the onset and offset locations of the target in polar coordinates. As shown in **Fig. 4B** for a representative subject, saccades were well accounted by both the onset (representative subject r^2^ = 0.83, mean ± s.e.m.; 0.81±0.01) and offset (representative subject r^2^ = 0.88, 0.87±0.01) target location, yet better by the latter than the former (paired t-test, t(10) = 14.74, p = 4.12 x 10^−8^; self-motion and no self-motion conditions considered jointly). In general, participants were reasonably accurate, but tended to undershoot the lateral location of targets (onset, slope; 0.77±0.03; offset; 0.79±0.03; onset vs. offset, paired t-test, t(10) = 7.44, p = 2.18 x 10^−5^). To ascertain whether humans saccade to the initial target location or incorporate knowledge of object-motion and saccade to its offset location, we fit a regression model (*y* ~ 1 + *β*_1_*onset* + *β*_2_*offset*) with *β*_1_ and *β*_2_ respectively weighting the target onset and offset locations. As shown in **Fig. 4C**, subjects placed more weight on the target offset than onset locations (onset vs. offset, no self-motion and self-motion, paired t-test, both t > 2.21 and p < 0.04). Similarly, while the offset regressor was significant for 11 (no self-motion) and 8 (self-motion) of the 11 subjects, the onset regressor was significant for 9 (no self-motion) and 4 (self-motion). Finally, given (i) our interest in comparing between a concurrent object- and self-motion condition requiring CI, and an object-motion only condition not requiring CI, and (ii) the above evidence that saccade behavior was mostly dictated by the target’s offset location, we binned the latter and examined whether gaze to target offset location was modulated by self-motion. Results showed no impact of self-motion on the landing saccade position (**Fig. 4D**, unpaired ANOVA, interaction term, p = 0.95). Thus, subjects’ initial inference of target position, as reflected by saccades, does not appear to be affected by misattribution of retinal velocity to object motion vs. self-motion.

Lastly, we examined the smooth pursuit that occurred after saccade offset. That is, for each trial we epoched and baseline-corrected the 500ms of eye movements following saccade offset. We split trials as a function of target velocity and self-motion condition. Strikingly, as shown in **Fig. 4E**(means across all subjects), while smooth pursuit was modulated by target velocity in the no self-motion condition, it was not during concurrent self- and object-motion. We quantify this effect by computing the mean velocity of eye displacement across the 500ms following saccade offset. This analysis reveals no dependence of eye velocity on target velocity in the self-motion condition (one-way ANOVA, F(10, 110) = 0.41, p = 0.93), and the presence of a strong modulation when object-motion was presented alone (one-way ANOVA, F(10, 110) = 21.7, p = 6.66×10^−22^, **Fig. 4F**). In fact, the gaze velocity along azimuth matched the target velocity in the absence of self-motion (one-way ANOVA on the difference between target and gaze velocity, F(10, 110) = 0.75, p = 0.67).

Together, the pattern of eye movements shows largely accurate saccading to the target offset regardless of the self-motion condition, followed by smooth pursuit of the target that is absent in the case of concurrent self- and object-motion, in contrast to the object-motion only condition and our prior work (where the target never moved; [22, 25]). That is, during the no self-motion condition gaze velocity matches that of targets, while it is null after concurrent self- and object-motion. Crucially, during smooth pursuit, the target is invisible in both cases, such that this difference is not simply a visually-driven effect.

## 3. Discussion

A wide array of studies have argued that human perception is scaffolded on the process of CI: we first hypothesize a generative structure (i.e., how “data” is generated by the world), and then use this hypothesis in perceiving [2, 5–10, 14–16, 30–37]. As such, some [3] have argued that CI is a unifying theory of brain function. Here, our main contribution is the derivation of CI predictions for a naturalistic and continuous-time task, and the demonstration of these signatures during closed-loop active sensing. Further, we demonstrate how using continuous-time tasks may allow us to index the temporal unfolding of computations that mediate CI.

We take the example of concurrent self- and object-motion where observers have to infer whether flow vectors on their retina are generated entirely by self-motion, by object-motion, or by some combination of the two. Empirically, we show that, in line with the derived predictions, humans are more likely to report moving targets as stationary at high velocities during concurrent self-motion. Similarly, during concurrent self- and object-motion, observers navigate to locations closer to the initial target location, as if the target were not moving. This effect is most pronounced when targets move at intermediate velocities: those that cannot be easily ascribed to a single interpretation, either self- or object-motion. We further support the claim that observers misattribute flow vectors caused by object-motion to self-motion by testing and corroborating two additional qualitative predictions. First, we note that lateral flow vectors at the focus of expansion are essentially null when moving forward and toward the focus of expansion (see inset in **Fig. 2A**). Thus, in the current setup there should be little to no misattribution of flow vectors when targets are presented centrally (i.e., near the focus of expansion), and this prediction was confirmed (**Fig. 3B**). Second, we highlight that our estimates of self-motion are not solely derived from optic-flow, but also from efference copies of motor commands, among other signals. Thus, we ran a second experiment in which the self-motion experienced during object-motion was not under the subjects’ control. We argue that passive self-motion, by virtue of lacking efference copies of motor commands that are present during active self-motion, should result in a less certain estimate of self-motion and thus result in a greater likelihood of misattributions. This prediction was also confirmed (**Fig. 3D**). Together, these findings strongly suggest that observers perform CI when navigating in the presence of objects that may or may not move.

The third and final experimental finding allowed us to gain some insight into the timecourse of the processes constituting CI. That is, we show that saccades to hidden targets already incorporate knowledge of the object velocity, with target offset position accounting for a larger fraction of variance than target onset position. The fact that these saccades to target were largely accurate may suggest that, at the time of saccade onset (~200-300ms from target onset), object-motion was appropriately parsed from self-motion. Prior studies [11] have shown CI during saccades, but these studies use saccades as a reporting mechanism, as opposed to allowing observers to use saccades in a natural and continuous-time environment. Next, we show that smooth gaze pursuit of the target following the initial saccade was strikingly different between the self-motion and no self-motion conditions. While in the no self-motion condition gaze velocity showed a gradient consistent with the target velocity, there was no modulation of gaze velocity by target velocity in the self-motion condition. This difference cannot be attributed to differences in optic flow, as at the time of smooth pursuit (~400-900ms after target onset) participants were moving and experiencing optic flow in both conditions. These results suggest that at the time of smooth pursuit, flow vectors caused by object- and self-motion were not properly parsed. Speculatively, these findings (i.e., parsing at the time of saccades but not at the time of smooth pursuit) are evocative of a cascade of events observed in neural data [4, 14–16] wherein primary sensory cortices segregate cues, later “associative” areas always integrate cues, and finally “higher-order” regions perform flexible CI. Interestingly, the established neural cascade occurs earlier than the herein described behavioral one (see c.f. **Fig. 4F** in [17]; neurally, segregation occurs between 0-100ms post-stimulus onset, forced integration occurs between 100 and 350ms, and CI thereafter).

We must mention a number of limitations in the current work. While our aim is to study CI within dynamic action-perception loops, much of our analysis relied on specific events frozen in time, and particularly at the end of trials (e.g., trajectory end-points and explicit reports). This was somewhat mitigated by the eye movement analyses and is a reflection of i) challenges associated with jointly modeling CI, path integration, and control dynamics, as well as ii) the fact that within the current paradigm inference over object-motion occurs only once, during the observation period. Namely, in future work we hope to leverage the full trajectories generated by participants in attempting to intercept moving, hidden targets. This, however, will require accounting for both idiosyncrasies emanating from path integration (e.g., putatively a slow-speed prior and cost functions that evolve with time and distance travelled [21, 23]), as well as derivation of optimal control policies (see [38–40] for recent attempts to model continuous behavior as rational, and then invert this model to deduce the dynamics of internal models). Similarly, at risk of losing generalizability, the modelling approach could be expanded to explicitly take flow vectors as input. More generally, however, these next steps in our modeling approach are designed to account for evolving beliefs, and thus require just that, behavior reflecting a protracted unfolding of an evolving belief. Examination of steering trajectories (**Fig. 2C**) did not suggest frequent and abrupt re-routings, as one would expect to occur during changing interpretations of the target motion (e.g., from moving to stationary). Instead, it appears that within the current paradigm observers made a causal inference once, early in the trial and while the target was visible.

Altogether, we derive normative predictions for how observers ought to intercept moving and hidden targets by inferring (i) whether objects are independently moving in the world, and (ii) if so, with what velocity. We then support the conjecture that humans perform CI in attributing flow vectors on their retina to different elements (i.e., self and object) during navigation. Lastly, we show that when allowed to evolve naturally in time, behavior may demonstrate a protracted unfolding of the computations characterizing CI. In the past we have shown that macaques will intuitively track the ongoing location of moving and hidden targets [24]. Hence, the current results demonstrating signatures of CI in attempting to “catch” moving targets opens the avenue for future studies of naturalistic and closed-loop CI at the single cell level.

## 4. Methods

### 4.1. Participants

Eleven participants (age range = 21-35 years old; 5 females) took part in the study. Participants had normal or corrected-to-normal vision, normal hearing, and no history of neurological or musculoskeletal disorders. The experimental protocol was approved by the University Committee on Activities Involving Human Subjects at New York University.

### 4.2. Experimental Materials and Procedure

Participants were tasked with virtually navigating to and stopping at the location of a briefly presented target (i.e., the “firefly”) via an analog joystick with two degrees of freedom (linear and angular velocity; CTI Electronics; Ronkonkoma, USA). Visual stimuli were rendered via custom code in Unity (Unity Technologies; San Francisco, USA) and displayed on a virtual reality (VR) head-mounted display with a built-in eye tracker (90 Hz; HTC VIVE Pro; New Taipei, Taiwan). The subjective vantage point was set to a height of 100cm above the ground, and participants had a field of view of 110° of visual angle.

The virtual scene was comprised of ground plane textural elements (isosceles triangles, base × height = 8.5 × 18.5 cm) that were randomly positioned and reoriented at the end of their lifetime (lifetime = 250 ms; floor density = 1.0 elements per 1 m^2^). On each trial, subjects were re-positioned at the center of the virtual world, and a target (a black circle of radius 30cm, luminance and color matched to ground plane triangles) was displayed at a radial distance of 300cm, and within a lateral range spanning from −10° to +10° (uniform distribution; 0° being defined as straight-ahead). On 10% of trials, the target was visible throughout the duration of the trial. On the rest, targets were presented for either 75ms or 150ms (equal probability). On 25% of trials, the target did not move (i.e., object velocity = 0cm/s). On the remaining 75% of trials, the target moved laterally at 3, 6, 10, 20, or 40cm/s, either leftward or rightward (equal probability). Target presentation time, velocity, and direction were randomly interleaved across trials and within blocks. Participants navigated toward the target with a maximum linear velocity (*v*_max_) of 200cm/s, and a maximum angular velocity (*w*_max_) of 90°/s. On every trial, after participants stopped to indicate their perceived location of the (hidden) target, they were asked to explicitly report whether the target had moved or not, by deflecting the joystick leftward (target did not move) or rightward (target moved). Participants were informed that their responses were logged via an auditory tone. No feedback on performance was given. Inter-trial intervals were not defined to participants (i.e., ground plane elements were always visible and continuous with the previous frame) and of random duration (uniform) between 300 and 600ms.

Every participant took part in 2 experiments, on separate days (each session lasting ~1 hour). In the first experiment, participants performed two blocks of 200 trials, in which they were always under full closed-loop conditions (i.e., motor output dictated sensory input). In one block, targets were presented given that participants were not moving (linear velocity < 1cm/s; no self-motion condition). In the other block, targets were presented given that participants were moving (linear velocity > 20cm/s; self-motion condition). Block order was randomized across subjects and participants were informed prior to each block whether they had to move (“self-motion” condition) or remain static (“no self-motion” condition) for targets to appear. In the second experiment, participants performed 4 blocks of 200 trials. At difference from the first experiment, here linear velocity was set (open-loop, under experimental and not subject control) during the target presentation. Linear velocity was either null (0cm/s; no self-motion condition), or had a Gaussian profile with peak amplitude equal to 200cm/s (self-motion condition). When participants were passively displaced, the linear perturbation lasted 1 second, and the target was presented during the peak of the perturbation (200cm/s) such that it would be at a distance of 300cm. After the perturbation, participants gained full access to the control dynamics (as in Experiment 1). Self-motion and no self-motion conditions were interleaved within a single block in Experiment 2. In total each participant completed 1200 trials.

### 4.3. Modeling

We here provide a basic outline of the normative model. A more detailed description is provided in the supplementary information. At the beginning of each trial, we assume the target to be visible for some observation time *T* = *N δt*, where *δt* is the duration of individual, discrete time steps in which our model is formulated. These time steps can be thought of as the time between individual video frames. While our results are independent of the specific choice of *δt*, discretization simplifies the model description and derivation of its properties. In the *n*th time step the target’s true location is *z*_*n*_ = *z*_*N*_ − (*N* − *n*)*vδt*, where *z*_*N*_ is the target’s true location at the end of the observation period, and where *v* is the target’s velocity. The target’s initial location *z*_1_ is assumed to be drawn from a uniform distribution over a wide range of *z*_1_’s. Its velocity is *v* = 0 (stationary target, *γ* = 0) with probability 1 − *p*_*γ*_ and otherwise (moving target, *γ* = 1) drawn from a normal distribution 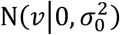 with mean zero and variance 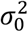 (a standard slow-velocity prior, [25, 26]). The observer has noisy observations, *x*_*n*_|*z*_*n*_ ~ N(*z*_*n*_, *σ*^2^/*δt*) whose variance is scaled by 1/*δt* to keep the results invariant to the choice of *δt* (i.e., choosing a smaller *δt* provides more, but individually less informative observations per unit time). Based on all observations, *x*_1:*N*_ ≡ *x*_1_,…, *x*_*N*_, the model estimates the probability of the target being stationary, *p*(*γ* = 0|*x*_1:*N*_), and the target’s velocity *p*(*v*|*x*_1:*N*_, *γ* = 1) in case it is moving. The first probability is used for its stationary reports (**Fig. 1B**). Both are used to decide on the model’s steering trajectory (**Fig. 1C**).

A lengthy derivation that is provided in the supplementary information yields the required posteriors. The resulting expressions are rather lengthy and thus not detailed here. We assume that the model reports that the target is stationary if *p*(*γ* = 0|*x*_1:N_) > *p*(*γ* = 1|*x*_1:N_), that is, if 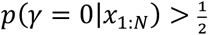, which is the case if

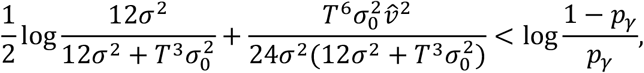

where 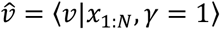 is the mean estimate of the target’s velocity under the assumption that it is moving. This decision rule imposes a threshold on this velocity estimate that depends on observation noise magnitude *σ*^"^ and observation time *T*, and leads to the stationary reports shown in **Fig. 1B**.

The model chooses the optimal steering angle and distance to maximize the probability of intercepting the target. To do so, we assume it uses the self-motion estimation model from Lakshminarasimhan et al. (2018) [19] in which the uncertainty (here measured as the variance of a Gaussian posterior over location) grows as *k*^2^*d*^2λ^, where *d* is the distance traveled, and *k* and *λ* are model parameters. For a given angle *θ* and travel time *t* (at fixed velocity *v*_*a*_) this provides a Gaussian posterior *p*(***z***_*a*_|*t*, *θ*) over 2D self-location ***z***_*a*_. Furthermore, assuming a known initial target distance and a lateral target motion, the model provides a similar posterior *p*(***z***_*o*_|*x*_1:N_, *t*) over 2D target location at time *t*, given by a weighted mixture of two Gaussians. The model then chooses *t*^*^ and *θ*^*^ that maximize *p*(***z***_*a*_ = ***z***_*o*_|*x*_1:N_, *t*, *θ*), that is, the likelihood of ending up at the target’s location when moving for some time *t*^*^ at angle *θ*^*^ (see supplementary information for respective expressions). Unfortunately, this maximization cannot be performed analytically, and thus we find the maximum by discretizing *t* and *θ*.

The model parameters where hand-tuned to provide a qualitative match to the human data. However, it is worth noting that the qualitative model behavior described in the main text is generic (i.e., fine-tuning is not necessary). Due to the simple nature of the model, and the simplified steering (i.e., control) model that only moved along a straight line, we did not attempt to achieve a quantitative match. The target was assumed to be at a fixed, known distance of 4m during the target observation period, and the observer moved at a constant *v*_*a*_ = 2m/s thereafter. A-priori, the target was assumed to be moving with *p*_*γ*_ = 0.3 and the standard deviation of its slow-velocity prior was *σ*_*0*_ = 0.5m/s. Observation times were set to either *T* = 0.075s or *T* = 0.150s, matching those of the experiment. Observation noise was set to *σ* = 0.0004m (no self-motion) or *σ* = 0.001m (self-motion). The path integration uncertainty parameters were set to *k* = 0.5 and *λ* = 0.5 (Wiener diffusion).

## Acknowledgments

This work was supported by NIH U19NS118246 (to GCD, DEA, and JD).

## Causal Inference Within Close Action-Perception Loops Supplementary Information

We here describe in detail the normative model that we used in the main text to predict behavioral signatures of causal inference.

### 1. TASK SETUP AND GENERATIVE MODEL

A visual target appears on the screen for *T* = *Nδt* seconds, or *N* time steps of size *δt*. While, for intuition, one can think of these time steps as individual video frames, our formulation is independent of the choice of *δt*. The target appears at location *z*_1_ and thereafter moves with velocity *v*. Therefore, its location in the *n*th time-step is given by

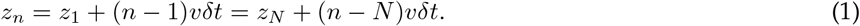

Across trials, we assume the target to be moving (*γ* = 1) with probability *p*(*γ* = 1) = *p_γ_*, and to be stationary (*γ* = 0) otherwise. When moving, its velocity is drawn from 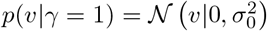, a normal distribution with zero mean and variance 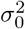. Overall, this leads to the prior over *υ* to be given by

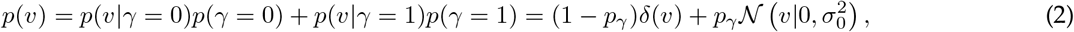

where *δ*(·) is the Dirac delta function. This is our causal inference prior as it encapsulates the two different hypotheses (stationary vs. moving target, *γ* = 0 vs. *γ* = 1) for what caused the sensory percepts, together with their associated priors on the underlying latent target velocity. We assume a uniform prior over the target’s initial location *z*_1_ over a bounded range, whose form we make more precise later.

In each time step, the observer makes a noisy observation *x_n_* of the target’s true location *z_n_*, distributed independently across time as

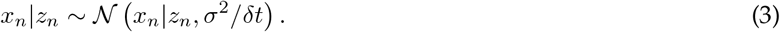

We here scale the observation’s variance by 1/*δt*, such that the overall amount of information that the observer receives per unit time remains invariant to the choice of *δt*.

Having observed *x*_1:*N*_ ≡ *x*_1_,…, *x*_*N*_, the observer wants to infer whether the target is moving or not, that is *p* (*γ* = 1|*x*_1:*N*_). Furthermore, they want to infer the target’s final location for a stationary target, *p* (*z_N_ γ* = 0|,*x*_1:*N*_), or the target’s final location and velocity for a moving target, *p* (*z_N_, υ|γ* = 1, *x*_1:*N*_). In the next section we derive these quantities. Following this, we turn to the question of how the observer uses these quantities to act upon them by reporting whether the target is stationary or moving, and how they steer toward the target.

### 2. INFERRING THE TARGET LOCATION AND VELOCITY

Here, we first start with assuming that the target is stationary to find *p* (*z_N_* |*γ* = 0, *x*_1:*N*_). Then, we assume a moving target and compute *p* (*z_N_, v*|*γ* = 1, *x*_1:*N*_). Lastly, we use the found expressions to derive *p* (*γ* = 1|*x*_1:*N*_).

#### 2.1. A stationary target

*γ* = 0.For the stationary case, we only need to find the posterior over the target’s single location *z* as its velocity is fixed to zero. Then, it is easy to show that

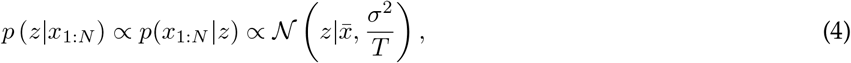

where we have implicitly conditioned on *γ* = 0, have assumed a uniform prior over *z* over the relevant range of *z*’s, and have defined

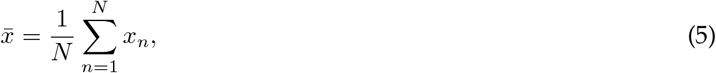

that is, the average observed location.

Causal inference also requires the marginal likelihood of ***x***_1:*N*_, which is given by

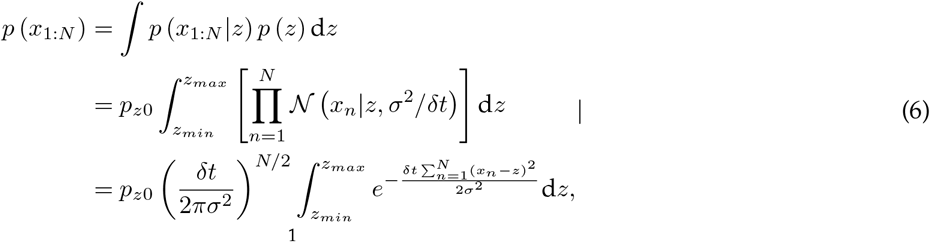

where we assumed a uniform prior of probability *p*(*z*) = *p_z_*_0_ over a wide *z*-range from *z_min_* to *z_max_*. The integral evaluates to

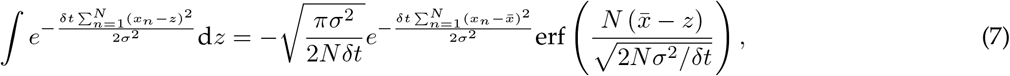

where the sum in the exponential equals *N* var (*x*), that is *N* times the empirical variance of *x*_1:*N*_. Furthermore, the error function approaches −1 for large *z_max_* and 1 for small *z_min_*, such that its contribution in the definite integral approaches −2. Therefore, the final marginal likelihood is

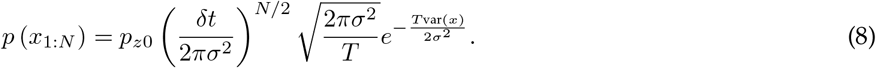

We can find the same result by writing down Bayes’ rule for *p*(*z|x*_1:*N*_) and solving for *p*(*x*_1:*N*_), which appears in the denominator.

#### 2.2 A moving target

*γ* = 1.Using the previous identity, *z_n_* = *z_N_* − (*N* − *n*)*υδt*, leads to the likelihood of each *x_n_* to be given by

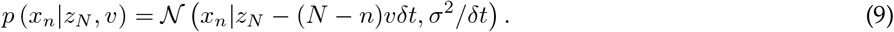

Our aim is to find the joint posterior over *z_N_* and *υ*, which is given by the expression

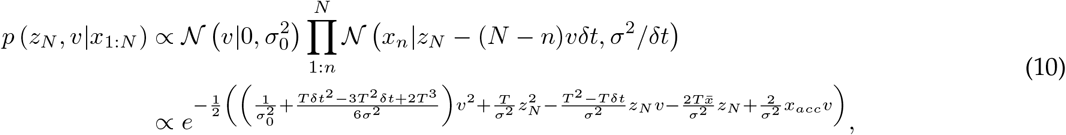

where we implicitly conditioned on *γ* = 1, have used the same definition of 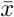 as further above, and have defined

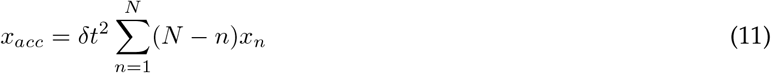

To find the posterior moments we first take *δt* → 0, removing all the *δt*-dependent terms. To de scribe th e full posterior, we denote it by

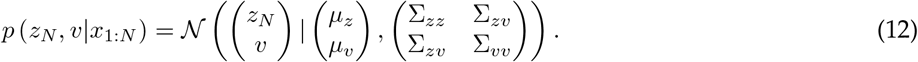

A lengthy, but unspectacular, derivation reveals

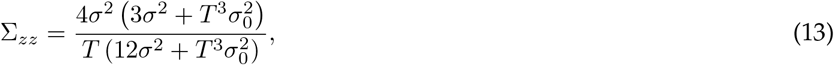

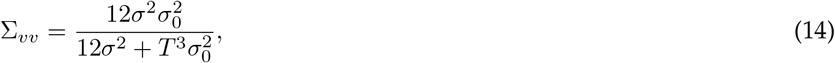

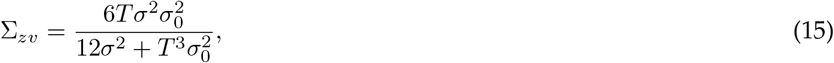

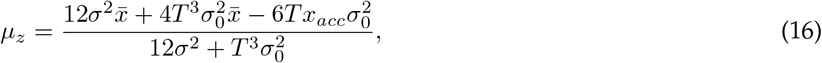

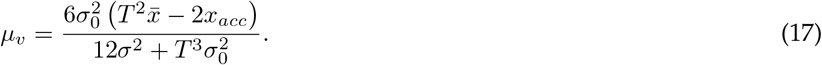

Interestingly, in the *σ*_0_ → ∞ limit, the posterior variance Σ_*zz*_ scales as 1/*T*, as before. The posterior variance Σ_*vv*_, in contrast, drops more rapidly with 1/*T* ^3^. Therefore, temporal integration of evidence provides qualitatively more velocity than location information. Intuitively, this is because any (*x_i_, x_j_*) pair can be used to infer velocity, whereas the location estimate relies on the across-*x_i_* average.

Understanding what the posterior means needs more work. In particular, let us define

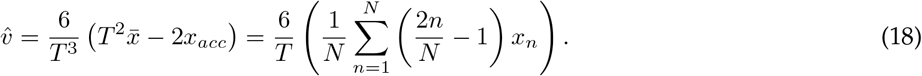

The term in (outer) parenthesis is a weighted sum of the *x_n_*’s. In this sum, *x*_1_ is weighted by −1, and *x_N_* by 1. Inbetween these extremes, the weights increase linearly from −1 to 1. Therefore, if we group equally-weighted terms (modulus the sign of the weight), the sum is a weighted combination of location differences, with the highest weight on *x_N_ − x*_1_, less weight on *x_N_*_−1_ – *x*_2_, and so on. To see how this can act as a velocity estimate, assume a noise-free *x_n_* = *vnδt*. Substituting this in the above expression shows that 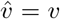 (this is where the 6/*T* pre-factor comes from). This justifies denoting it 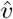.

Substituting 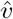 into the posterior means results in the new expressions

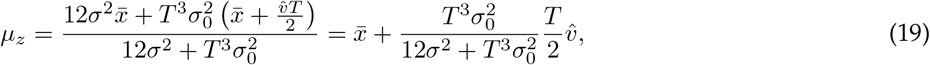

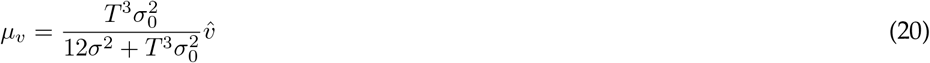

This shows that the posterior mean *μ_z_* start with 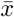 for small *T*, and then shifts toward 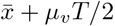, which is the mean location plus half the estimated distance that the target moved, which is sensible. The posterior mean *μ_v_* is initially biased toward zero, due to the prior, and later approaches 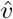. Overall, with 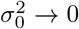, the posterior approaches that for a stationary target, as desired.

To find the marginal likelihood, we solve Bayes’s rule for the posterior for *p*(*x*_1:*N*_), which yields

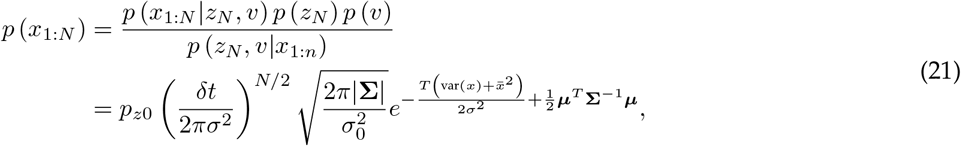

where we have chosen *z_N_* = *v* = 0 for the second equality (as the expression holds for any choice of *z_N_* and *υ*), and ***μ*** and **Σ** denote the posterior mean and covariance. The remaining terms evaluate to

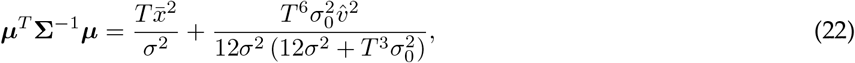

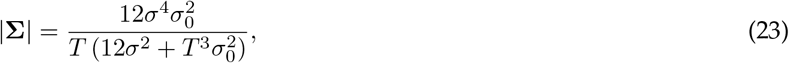

such that the marginal likelihood becomes

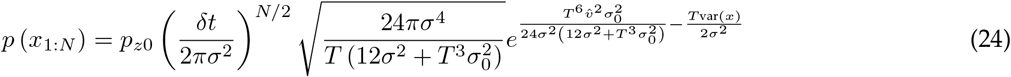

As for the posterior, this marginal likelihood approaches that for a stationary target with 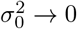.

#### 2.3 Is the target stationary or moving?

To find the full posterior over the target’s state, we use the causal inference target velocity prior, 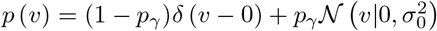, with which the posterior becomes

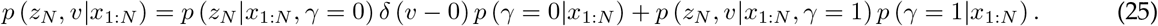

In this mixture distribution, the first mixture component is that for the stationary target, and the second that for the moving one. These two components are weighted by the causal modeling posterior *p*(*γ|x*_1:*N*_) which indicates the probability of the target being stationary or moving given the data. This probability can again be found by Bayes’ rule, and results in

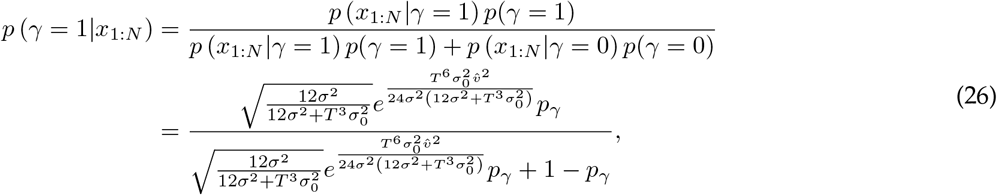

where the last expression results from substituting the marginal likelihoods for the stationary and moving target, and cancelling all the shared terms. This expression shows that, for a uniform prior, *p*(*γ* = 0) = *p*(*γ* = 1) = 1/2, the target is deemed more likely moving for larger velocity estimates 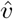. If this estimate is zero, that is, 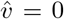, then, the more time has passed, the less likely is the target considered to be moving.

### 3. ACTING UPON THE INFERRED TARGET LOCATION AND VELOCITY

#### 3.1. Choosing stationary vs. moving

Decision-makers would choose between a stationary and a moving target according to *p* (*γ*|*x*_1:*N*_). In particular, they would decide that the target is moving if *p*(*γ* = 1|*x*_1:*N*_) > 1/2, that is, if

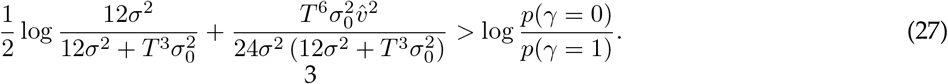

##### 3.1.1. Empirical distribution of stationary reports

As experimenters we cannot directly observe 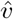, such that we need to estimate it. Furthermore, it will fluctuate across trials, even if the same evidence is presented, making the decisions more noisy.

To build a model for 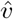 we note that 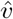 is a weighted sum of *x_n_*’s, and rely on our generative assumptions of *x_n_* for a stationary and a moving target. For a stationary target, we have *x_n_ ∽ *N* (z, σ*^2^/*δt*). In this case, 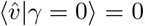, and its variance is given by

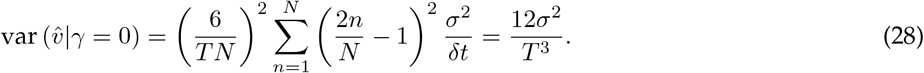

This variance decreases rapidly with time, as more and more *x_n_*’s are used to estimate 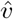. This results in the required moments

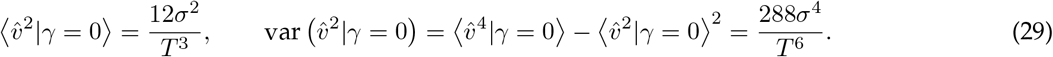

If we denote the 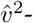-related term in the above decision criterion, Eq. (27), by *α*, this *α* has moments

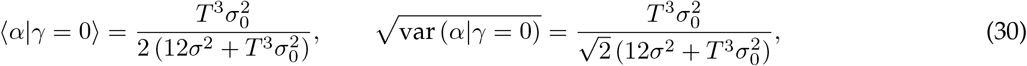

leading to the signal-to-noise ratio 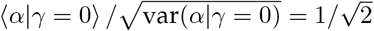. More relevant, under the assumption that *α* is Gaussian whose parameters are fully determined by mean and variance, the probability that the decision criterion is met becomes

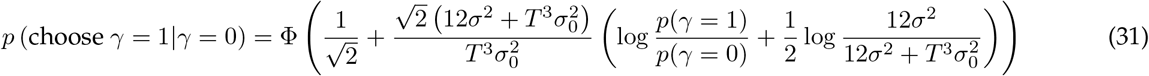

For *T* → 0 or 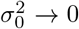 this probability is dominated by the prior and lead to a choice of *γ* = 1 if *p*(*γ* = 1) >*p*(*γ* = 0). It shrinks with increasing *T*, as more evidence results in higher certainty that the target is not moving. Increasing the observation noise *σ*^2^ has two counteracting effects. First, it increases the pre-factor to the inner-most brackets, thus boosting the prior. Second it results in a weaker drop of the last term in brackets with time, indicating that more evidence will be required to discard the possibility that the target is moving.

For a moving target, *x_n_* ~ *N* (*z_N_* − (*N* − *n*)*vδt, σ*^2^/*δt*). This yields 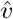 to be Gaussian, with moments

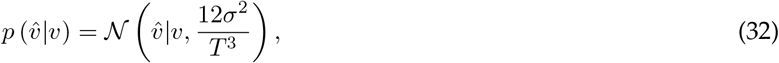

which only differs in the non-zero mean from the 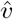 for the stationary case. With the above, the moments of 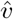 are given by

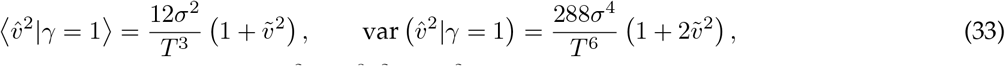

where we have defined the time-rescaled velocity 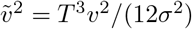. The previously defined *α* then has moments

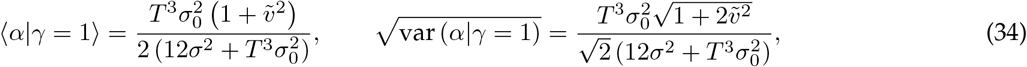

leading to the signal-to-noise ratio 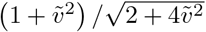 which approaches the linear function 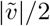 for larger 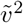. Plugging these moments into the decision criteria and assuming Gaussianity leads to the choice probability

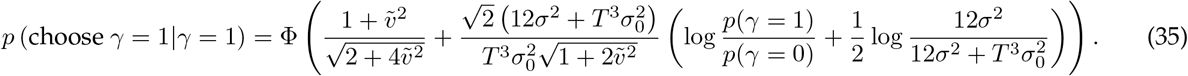

For 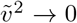, this probability becomes equivalent to the one for *γ* = 0. The larger 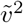, the stronger the influence of the first term, and the weaker the influence of the remaining terms. In particular, the larger 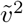, the higher the probability of choosing *γ* = 1.

A non-approximate approach to computing *p* (choose *γ* = 1 *γ* = 1) is to re-write the decision criterion for *p* (*γ* = 1|*x*_1:*N*_) > 1/2 as

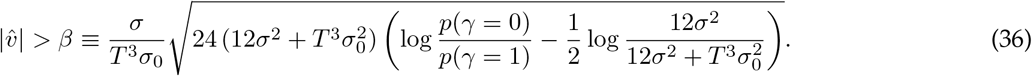

This criterion is only valid if the term in square-roots is non-negative, which is guaranteed as long as

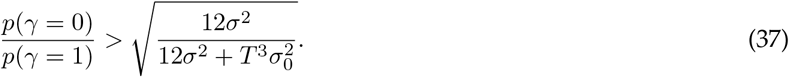

This is always satisfied if *p*(*γ* = 0) >*p*(*γ* = 1). In general, it requires little evidence about the target’s motion (i.e., small *T* and 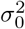 and large *σ*^2^), and a relatively strong prior toward the target not moving. If the above condition is violated, the decision criterion becomes 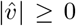, which is always satisfied. That is, under these circumstances, the target will always be considered moving.

Assuming there is a non-zero chance of the target being stationary, then the probability of the decision-maker reporting a stationary target depends on the perceived 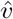 which is Gaussian in the true *υ* (see above). Then, as *β* ≥ 0, 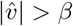 is satisfied if either 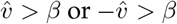. As these two options are mutually exclusive, their joint probability sums and is given by

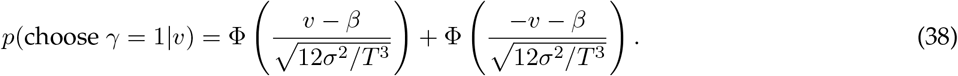

Simulations confirmed that these expressions match simulated choices.

### 4. STAGE II: INTERCEPTING THE TARGET

We assume agents travel at constant velocity, such that we only need to determine travel direction and stopping time for the best target interception. The objective is to minimize the expected cost, ⟨*c* (*z_a_*(*t*)⟩, *z_o_*(*t*)), where *z_a_* is the agent’s location, *z_o_* is the target’s location, and the expectation is over the uncertainty involving both. We will assume a simple cost function *c* (*z_a_, z_o_*) = −*δ* (*z_a_ – z_o_*), which is minimized if *z_a_* = *z_o_*.

The task itself is two-dimensional: the target appears at a certain distance and can move only laterally. We will assume that the depth is known and denoted *z_o,d_* (*o* for object, and *d* for depth). The agent moves at a constant velocity *υ_a_* at angle *θ* (*θ* = 0 is straight-ahead) for some time *t*. Then, what needs to be determined for optimal interception is the agent’s stopping time *t*^*^ and the angle *θ*^*^ that minimizes the expected cost,

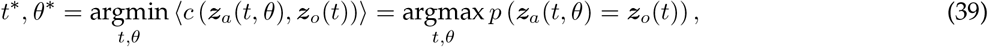

where the second equality follows from the delta cost-function.

#### 4.1. Agent motion model

We will use the self-motion estimation model from Lakshminarasimhan et al. (2018), where they assume a Weber-like variance scaling of the self-location estimate,

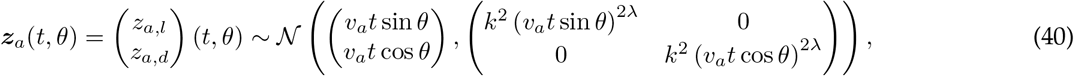

with parameters *k* (*k*^2^ is the variance scaling factor) and *λ* (determines the sub/supra-linearity of the variance scaling), and where we have assumed *z_a_*(0, *θ*) = (0, 0)^*T*^.

#### 4.2. Target motion model

As the target depth is known, we only need to estimate its initial lateral location and eventual velocity. To do so, we use the target location/velocity estimates derived further above, which provide the joint estimate

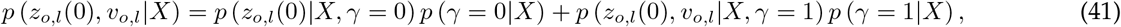

where *X ≡ x*_1:*N*_ are all target observations up to the target offset, *z_o,l_*(0) is the inferred lateral position at target offset (measured relative to agent), *v_o,l_* is the target’s lateral velocity, and *γ* ∈ {0, 1} denotes the target being stationary/moving. To compute the posterior over *z_o,l_*(*t*) for *γ* = 1, we use *z_o,l_*(*t*) = *z_o,l_*(0) + *v_o,l_* to get, as before,

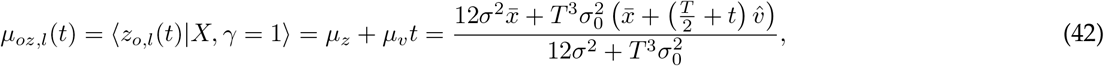

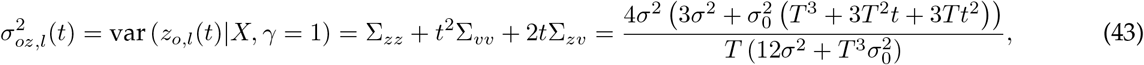

where *μ_z_*, *μ_υ_*, and the Σ’s are the posterior moments of *p*(*z_o,l_*(0), *v_o,l_|X, γ* = 1), Eq. (12). Overall, this leads to the posterior to be given by

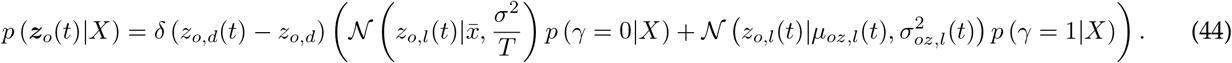

#### 4.3. Optimal steering angle and stopping time

To find the optimal steering angle *θ*^*^ and stopping time *t*^*^, we use the independence of the components of *z_a_*(*t*) and *z_o_*(*t*) to find

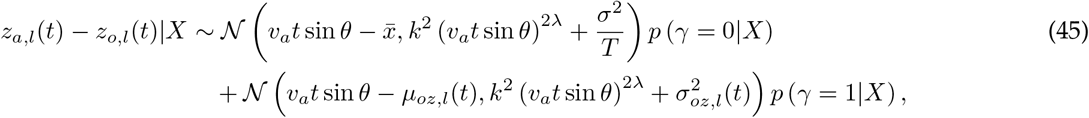

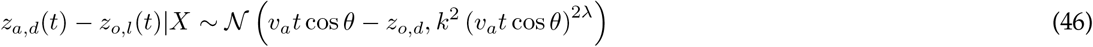

The optimal *θ*^*^ and *t*^*^ maximize the probability of both differences being zero. This leads to the expression

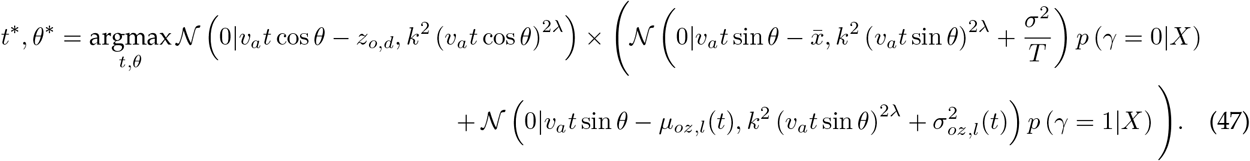

This optimization is complex and doesn’t have an analytical solution. Therefore, we need to use numerical optimization to find *θ*^*^ and *t*^*^.

For good initial guesses for this optimization, we consider the moving and stationary target case in isolation. For a stationary target, the Gaussian peaks at 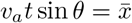 and *v_a_t* cos *θ* = *z_o,d_*, which leads to parameters

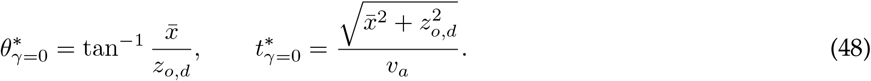

For a moving target, the Gaussian peaks at *υ_a_t* sin *θ* = *μ_oz,l_*(*t*) = *μ_z_* + *μ_υ_t* and *v_a_t* cos *θ* = *z_o,d_*, resulting in

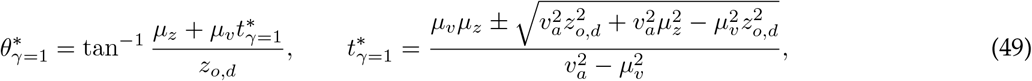

where we provide two solutions to 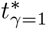 which result from solving a quadratic equation, and where *μ_z_* and *μ_v_* are the posterior moments of *p* (*z_o,l_*(0), *v_o,l_*|*X, γ* = 1). To identify a unique 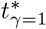 we assume that 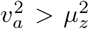, such that the agent is guaranteed to be able to catch up with the target. Furthermore, we require 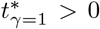 which is guaranteed to be violated if 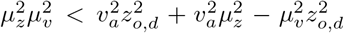, or, equally, 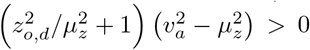. The latter holds if both 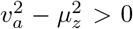 (guaranteed by assumption) and 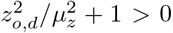, or 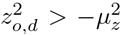. As the right-hand side of the latter is always negative, the last inequality is always true, confirming that the only solution to 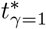 is the one with a sum (rather than difference) in the numerator. Substituting the expressions of *μ_z_* and *μ_v_* in terms of 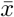 and 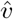 does not lead to any appreciable simplifications.

Together, these two solutions allow us to approximate the optimal heading directions and stopping times by

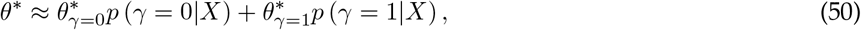

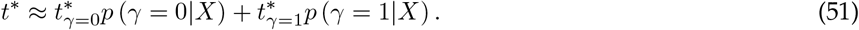

We use these approximations to initiate a gradient-descent procedure to find the correct *t* ^*^ and *θ*^*^.

#### 4.4. Experimentally observable optimal stopping, and perceived velocity estimate

As for the stationarity reports, the experimenter does not observe 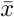 and 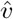 and needs to marginalize over them. They are distributed as

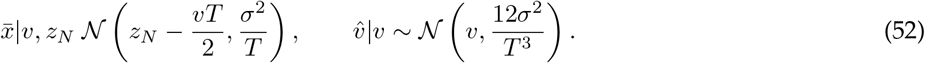

We estimate the associated statistics of *θ*^*^ and *t*^*^, by a Monte Carlo approximation. That is, we draw multiple 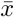 and 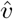, compute the associated *θ*^*^ and *t*^*^, and use those to compute the posterior distributions over *θ*^*^ and *t*^*^.

To assess the decision maker’s estimate of the target’s velocity, we will infer the assumed velocity from the decision maker’s stopping location. The actual stopping location could feature a mismatch between the decision maker’s depth and that of the target. By our initial assumption that the decision maker knows the target’s depth, this mismatch arises from the decision maker’s actual location while actively steering, and thus doesn’t refect the decision maker’s estimate. Thus, instead of using the agent’s final location as a measure for the assumed velocity, we will instead use the point *z_o,d_* tan *θ*^*^ and time 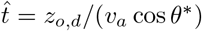 at which the agent crossed the target’s path (i.e. reaches depth *z*_0*,d*_).

Therefore, we won’t take the depth mismatch into account, and instead only focus on lateral location, using

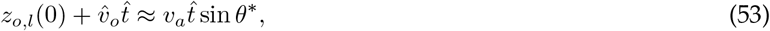

where the left-hand side is the target’s location at time 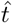(assuming velocity 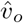), and the right-hand side is the decision maker’s location at the same time. Re-expressing the above in terms of 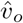 results in

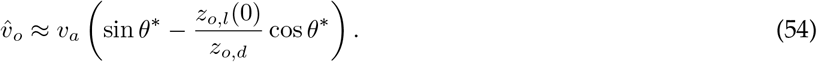

## References

Wolpert DM, Miall RC, Kawato M. Internal models in the cerebellum. Trends Cogn Sci. 1998 Sep 1;2(9):338–47. doi: 10.1016/s1364-6613(98)01221-2. PMID: 21227230.

Körding KP, Beierholm U, Ma WJ, Quartz S, Tenenbaum JB, Shams L. Causal inference in multisensory perception. PLoS One. 2007 Sep 26;2(9):e943. doi: 10.1371/journal.pone.0000943. PMID: 17895984; PMCID: PMC1978520.

Shams L, Beierholm U. Bayesian causal inference: A unifying neuroscience theory. Neurosci Biobehav Rev. 2022 Jun;137:104619. doi: 10.1016/j.neubiorev.2022.104619. Epub 2022 Mar 21. PMID: 35331819.

Noppeney U. Perceptual Inference, Learning, and Attention in a Multisensory World. Annu Rev Neurosci. 2021 Jul 8;44:449–473. doi: 10.1146/annurev-neuro-100120-085519. Epub 2021 Apr 21. PMID: 33882258.

Odegaard B, Wozny DR, Shams L. Biases in Visual, Auditory, and Audiovisual Perception of Space. PLoS Comput Biol. 2015 Dec 8;11(12):e1004649. doi: 10.1371/journal.pcbi.1004649. PMID: 26646312; PMCID: PMC4672909.

Acerbi L, Dokka K, Angelaki DE, Ma WJ. Bayesian comparison of explicit and implicit causal inference strategies in multisensory heading perception. PLoS Comput Biol. 2018 Jul 27;14(7):e1006110. doi: 10.1371/journal.pcbi.1006110. PMID: 30052625; PMCID: PMC6063401.

Samad M, Chung AJ, Shams L. Perception of body ownership is driven by Bayesian sensory inference. PLoS One. 2015 Feb 6;10(2):e0117178. doi: 10.1371/journal.pone.0117178. PMID: 25658822; PMCID: PMC4320053.

Noel JP, Samad M, Doxon A, Clark J, Keller S, Di Luca M. Peri-personal space as a prior in coupling visual and proprioceptive signals. Sci Rep. 2018 Oct 25;8(1):15819. doi: 10.1038/s41598-018-33961-3. PMID: 30361477; PMCID: PMC6202371.

Yang S, Bill J, Drugowitsch J, Gershman SJ. Human visual motion perception shows hallmarks of Bayesian structural inference. Scientific reports. 2021 Feb 12;11(1):1–4.

Bill J, Gershman SJ, Drugowitsch J. Visual motion perception as online hierarchical inference. Nature communications. 2022 Dec 1;13(1):1–7.

Mohl JT, Pearson JM, Groh JM. Monkeys and humans implement causal inference to simultaneously localize auditory and visual stimuli. J Neurophysiol. 2020 Sep 1;124(3):715–727. doi: 10.1152/jn.00046.2020. Epub 2020 Jul 29. PMID: 32727263; PMCID: PMC7509303.

Dokka K, Park H, Jansen M, DeAngelis GC, Angelaki DE. Causal inference accounts for heading perception in the presence of object motion. Proc Natl Acad Sci U S A. 2019 Apr 30;116(18):9060–9065. doi: 10.1073/pnas.1820373116. Epub 2019 Apr 17. PMID: 30996126; PMCID: PMC6500172.

Fang W, Li J, Qi G, Li S, Sigman M, Wang L. Statistical inference of body representation in the macaque brain. Proc Natl Acad Sci U S A. 2019 Oct 1;116(40):20151–20157. doi: 10.1073/pnas.1902334116. Epub 2019 Sep 3. PMID: 31481617; PMCID: PMC6778219.

Rohe T, Noppeney U. Cortical hierarchies perform Bayesian causal inference in multisensory perception. PLoS Biol. 2015 Feb 24;13(2):e1002073. doi: 10.1371/journal.pbio.1002073. Update in: PLoS Biol. 2021 Nov 18;19(11):e3001465. PMID: 25710328; PMCID: PMC4339735.

Rohe T, Noppeney U. Distinct Computational Principles Govern Multisensory Integration in Primary Sensory and Association Cortices. Curr Biol. 2016 Feb 22;26(4):509–14. doi: 10.1016/j.cub.2015.12.056. Epub 2016 Feb 4. PMID: 26853368.

Rohe T, Ehlis AC, Noppeney U. The neural dynamics of hierarchical Bayesian causal inference in multisensory perception. Nat Commun. 2019 Apr 23;10(1):1907. doi: 10.1038/s41467-019-09664-2. PMID: 31015423; PMCID: PMC6478901.

Aller M, Noppeney U. To integrate or not to integrate: Temporal dynamics of hierarchical Bayesian causal inference. PLoS Biol. 2019 Apr 2;17(4):e3000210. doi: 10.1371/journal.pbio.3000210.

Qi G., Fang W., Li S., Li J., Wang, L. (2021). Neural dynamics of causal inference in the macaque frontoparietal circuit. bioRxiv 2021.12.06.469042; doi: https://doi.org/10.1101/2021.12.06.469042

French RL, DeAngelis GC. Multisensory neural processing: from cue integration to causal inference. Curr Opin Physiol. 2020 Aug;16:8–13. doi: 10.1016/j.cophys.2020.04.004. Epub 2020 Apr 18. PMID: 32968701; PMCID: PMC7505234.

Rideaux R, Storrs KR, Maiello G, Welchman AE. How multisensory neurons solve causal inference. Proc Natl Acad Sci U S A. 2021 Aug 10;118(32):e2106235118. doi: 10.1073/pnas.2106235118. PMID: 34349023; PMCID: PMC8364184.

Lakshminarasimhan KJ, Petsalis M, Park H, DeAngelis GC, Pitkow X, Angelaki DE. A Dynamic Bayesian Observer Model Reveals Origins of Bias in Visual Path Integration. Neuron. 2018 Jul 11;99(1):194–206.e5. doi: 10.1016/j.neuron.2018.05.040. Epub 2018 Jun 21. PMID: 29937278; PMCID: PMC6190923.

Lakshminarasimhan KJ, Avila E, Neyhart E, DeAngelis GC, Pitkow X, Angelaki DE. Tracking the Mind’s Eye: Primate Gaze Behavior during Virtual Visuomotor Navigation Reflects Belief Dynamics. Neuron. 2020 May 20;106(4):662–674.e5. doi: 10.1016/j.neuron.2020.02.023. Epub 2020 Mar 13. PMID: 32171388; PMCID: PMC7323886.

Noel JP, Lakshminarasimhan KJ, Park H, Angelaki DE. Increased variability but intact integration during visual navigation in Autism Spectrum Disorder. Proc Natl Acad Sci U S A. 2020 May 19;117(20):11158–11166. doi: 10.1073/pnas.2000216117. Epub 2020 May 1. PMID: 32358192; PMCID: PMC7245105.

Noel JP, Caziot B, Bruni S, Fitzgerald NE, Avila E, Angelaki DE. Supporting generalization in non-human primate behavior by tapping into structural knowledge: Examples from sensorimotor mappings, inference, and decision-making. Prog Neurobiol. 2021 Jun;201:101996. doi: 10.1016/j.pneurobio.2021.101996. Epub 2021 Jan 14. PMID: 33454361; PMCID: PMC8096669.

Noel, J.P., Balzani, E., Avila, E., Lakshminarasimhan, K., Bruni, S., Alefantis, P., Savin, C., Angelaki. D. (2022). Coding of latent variables in sensory, parietal, and frontal cortices during virtual closed-loop navigation. eLife 11, e80280

Alefantis P, Lakshminarasimhan K, Avila E, Noel JP, Pitkow X, Angelaki DE. Sensory Evidence Accumulation Using Optic Flow in a Naturalistic Navigation Task. J Neurosci. 2022 Jul 6;42(27):5451–5462. doi: 10.1523/JNEUROSCI.2203-21.2022. Epub 2022 May 31. PMID: 35641186; PMCID: PMC9270913.

Stocker AA, Simoncelli EP (2006) Noise characteristics and prior expectations in human visual speed perception. Nat Neurosci 9:578–585. doi:10.1038/nn1669 pmid: 16547513

Weiss Y, Simoncelli EP, Adelson EH (2002) Motion illusions as optimal percepts. Nat Neurosci 5:598–604. doi: 10.1038/nn0602-858 pmid: 12021763

Ma WJ. Organizing probabilistic models of perception. Trends Cogn Sci. 2012 Oct;16(10):511–8. doi: 10.1016/j.tics.2012.08.010

Noel JP, Shivkumar S, Dokka K, Haefner RM, Angelaki DE. Aberrant causal inference and presence of a compensatory mechanism in autism spectrum disorder. Elife. 2022 May 17;11:e71866. doi: 10.7554/eLife.71866. PMID: 35579424; PMCID: PMC9170250.

Alais D, Burr D. The ventriloquist effect results from near-optimal bimodal integration. Curr Biol. 2004 Feb 3;14(3):257–62. doi: 10.1016/j.cub.2004.01.029. PMID: 14761661.

Wallace MT, Roberson GE, Hairston WD, Stein BE, Vaughan JW, Schirillo JA. Unifying multisensory signals across time and space. Exp Brain Res. 2004 Sep;158(2):252–8. doi: 10.1007/s00221-004-1899-9. Epub 2004 Apr 27. PMID: 15112119.

Wozny DR, Beierholm UR, Shams L. Probability matching as a computational strategy used in perception. PLoS Comput Biol. 2010 Aug 5;6(8):e1000871. doi: 10.1371/journal.pcbi.1000871. PMID: 20700493; PMCID: PMC2916852.

de Winkel KN, Katliar M, Diers D, Bülthoff HH. Causal Inference in the Perception of Verticality. Sci Rep. 2018 Apr 3;8(1):5483. doi: 10.1038/s41598-018-23838-w. PMID: 29615728; PMCID: PMC5882842.

Cao Y, Summerfield C, Park H, Giordano BL, Kayser C. Causal Inference in the Multisensory Brain. Neuron. 2019 Jun 5;102(5):1076–1087.e8. doi: 10.1016/j.neuron.2019.03.043. Epub 2019 Apr 29. PMID: 31047778.

Perdreau F, Cooke JRH, Koppen M, Medendorp WP. Causal inference for spatial constancy across whole body motion. J Neurophysiol. 2019 Jan 1;121(1):269–284. doi: 10.1152/jn.00473.2018. Epub 2018 Nov 21. PMID: 30461369.

Magnotti JF, Beauchamp MS. A Causal Inference Model Explains Perception of the McGurk Effect and Other Incongruent Audiovisual Speech. PLoS Comput Biol. 2017 Feb 16;13(2):e1005229. doi: 10.1371/journal.pcbi.1005229. PMID: 28207734; PMCID: PMC5312805.

Kumar A, Wu Z, Pitkow X, Schrater P. Belief dynamics extraction. Cogsci. 2019 Jul;2019:2058–2064. PMID: 33367289; PMCID: PMC7754614.

Kwon M, Daptardar S, Schrater P, Pitkow X. Inverse Rational Control with Partially Observable Continuous Nonlinear Dynamics. Adv Neural Inf Process Syst. 2020 Dec;33:7898–7909. PMID: 34712038; PMCID: PMC8549572.

Straub, D., Rothkopf, C.A. (2021). Putting perception into action: Inverse optimal control for continuous psychophysics; bioRxiv 2021.12.23.473976; doi: https://doi.org/10.1101/2021.12.23.473976

